# Optogenetically induced reward and ‘frustration’ memory in larval *Drosophila*

**DOI:** 10.1101/2022.07.20.500885

**Authors:** Juliane Thoener, Aliće Weiglein, Bertram Gerber, Michael Schleyer

**Affiliations:** Leibniz Institute for Neurobiology, Department of Genetics, Magdeburg, Germany; Institute of Anatomy, Otto von Guericke University Magdeburg, Germany; Institute of Biology, Otto von Guericke University Magdeburg, Germany; Center for Behavioral Brain Sciences, Magdeburg, Germany; Institute for the Advancement of Higher Education, Hokkaido University, Japan

**Keywords:** Timing-dependent valence reversal, associative learning, reinforcement, dopamine, mushroom body

## Abstract

Humans and animals alike form oppositely valenced memories for stimuli that predict the occurrence versus the termination of a reward: appetitive ‘reward’ memory for stimuli associated with the occurrence of a reward and aversive ‘frustration’ memory for stimuli that are associated with its termination. We characterize these memories in larval *Drosophila* using a combination of Pavlovian conditioning, optogenetic activation of the dopaminergic central-brain DAN-i1^864^ neuron, and high-resolution video-tracking. This reveals their dependency on the number of training trials and the duration of DAN-i1^864^ activation, their temporal stability, and the parameters of locomotion that are modulated during memory expression. Together with previous results on ‘punishment’ versus ‘relief’ learning by DAN-f1 neuron activation, this reveals a 2×2 matrix of timing-dependent memory valence for the occurrence/ termination of reward/ punishment. These findings should aid the understanding and modelling of how brains decipher the predictive, causal structure of events around a target reinforcing occurrence.

## Introduction

Pleasurable events induce positive affect when they occur, and negative affect when they terminate. Accordingly, humans and animals alike form appetitive memory for stimuli that are associated with the occurrence of reward (‘reward’ memory), and aversive memory for stimuli associated with its termination (‘frustration’ memory) (Hellstern et al., 1998; Solomon and Corbit, 1974) (the same is observed for painful events, with inverted signs: Gerber et al., 2014; Solomon and Corbit, 1974). Although the mechanisms of reward memory are understood in considerable detail (Schultz, 2015; Waddell, 2013), much less is known about frustration memory (Felsenberg et al., 2013; Hellstern et al., 1998). Such an incomplete picture of how pleasurable events are processed may lead both computational modelling of neural networks and the understanding of related pathology astray. In this context, we provide the first characterization of the learning from the occurrence and termination of a central-brain reward signal.

For such an endeavour, larval *Drosophila melanogaster* are an attractive study case. Robust paradigms of associative learning and resources for cell-specific transgene expression are available (Eschbach et al., 2020; Li et al., 2014; Saumweber et al., 2018). Moreover, their central nervous system is compact and consists of only approximately 10,000 neurons (Bossing et al., 1996; Larsen et al., 2009). This has allowed for the reconstruction of its chemical-synapse connectome, including the mushroom bodies as the associative memory centre of insects (Eichler et al., 2017). It has turned out that the circuits and mechanisms underlying associative learning in larvae largely parallel those of adults (Eschbach and Zlatic, 2020; Thum and Gerber, 2019; adult flies: Li et al., 2020; Takemura et al., 2017). Sensory projection neurons establish a sparse, combinatorial representation of the sensory environment across the mushroom body Kenyon cells (KCs). The KC axons are intersected by mostly dopaminergic modulatory neurons (DANs) and by mushroom body output neurons (MBONs), which eventually connect towards the motor system to adapt the parameters of locomotion (Eschbach et al., 2021; Paisios et al., 2017; Schleyer et al., 2020; Thane et al., 2019). Axonal regions of DANs and MBON dendritic regions are organized as compartments in which matched-up individual DANs and MBONs form local circuits with the KCs (**Fig. 1A**). Of the eight DANs innervating the mushroom body, two mediate rewarding effects and three mediate punishing effects (Eschbach et al., 2020; Saumweber et al., 2018). When an odour is encountered, and a reward or punishment signal reaches a given compartment, the strength of the local synapses from the odour-activated KCs to the MBON(s) of that same compartment is modified. As a result, the animal’s behaviour towards the odour in question is changed when the odour is encountered again.

**Fig. 1:**
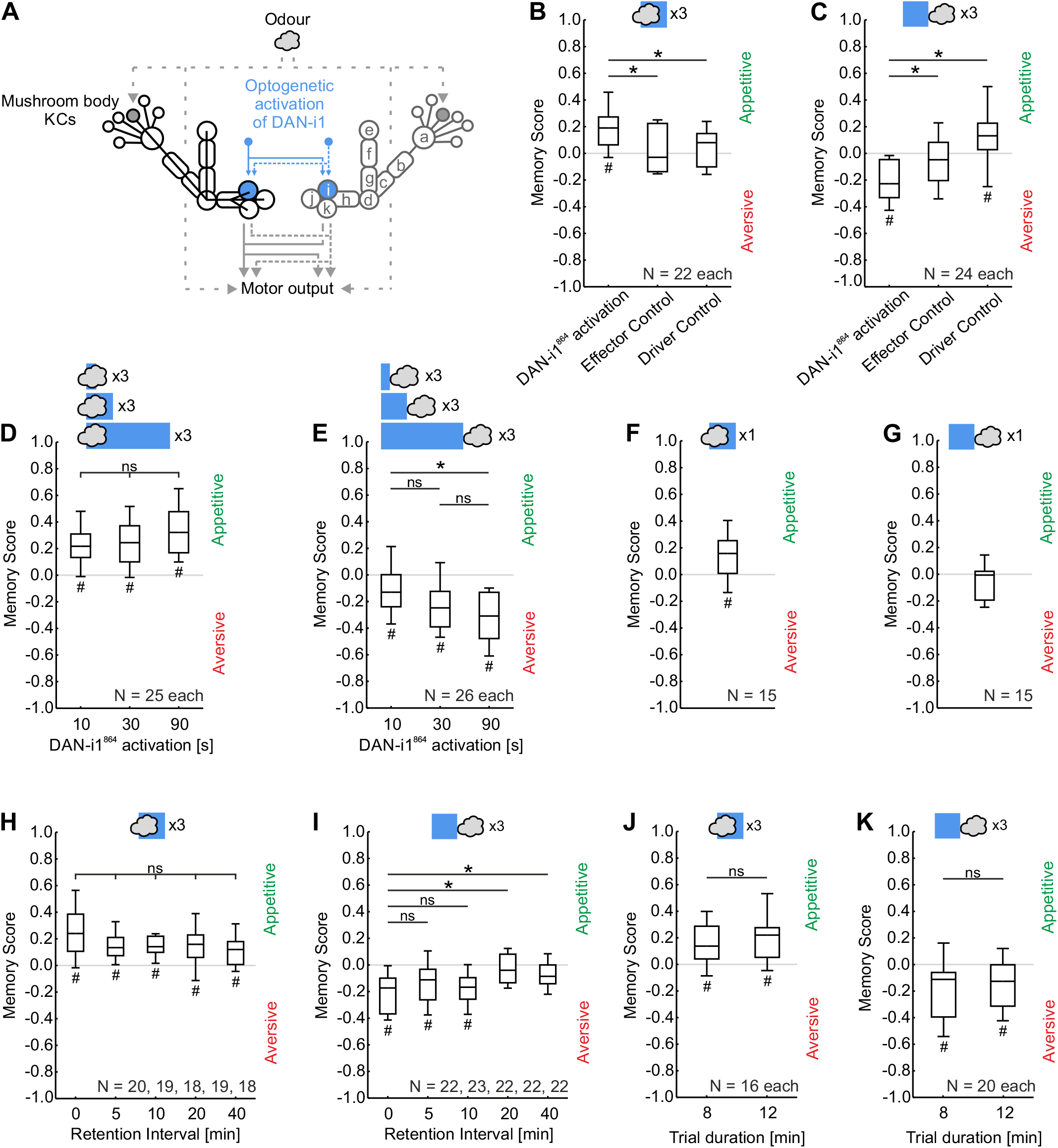
Reward and frustration memory by optogenetic DAN-i1^864^ activation. **(A)** Schematic of the DAN-i1 neuron (blue) innervating the i-compartment of the mushroom body. Kenyon cells (KCs) feature a higher-order odour representation. **(B, C)** Larvae were trained with three trials of odour (cloud) and DAN-i1^864^ activation (blue square) and subsequently tested for their odour preference. Positive and negative Memory Scores reflect appetitive or aversive associative memory, respectively (**Fig. S1**). **(B)** Upon forward training, only the experimental genotype showed appetitive memory. **(C)** Upon backward training, aversive memory was only shown by the experimental genotype. **(D, E)** Larvae received three trials of forward or backward training with varying durations of DAN-i1^864^ activation. **(D)** For forward training, odour presentation always preceded DAN-i1^864^ activation by 10 s. Appetitive memory of equal strength was observed in all cases. **(E)** For backward training, odour presentation always started at the offset of the DAN-i1^864^ activation. Increased durations of DAN-i1^864^ activation supported increased aversive memory. (**F, G**) With a single training trial, forward training established appetitive memory **(F),** whereas backward training established no aversive memory **(G)**. **(H, I)** When three training trials were conducted, larvae showed appetitive memory up to at least 40 min after forward training **(H)**, whereas aversive memory after backward training has decayed after just 20 min. Note that the training trial duration in these experiments was 8 min instead of 12 min as in **(B-G)**. The results in **(J, K)** show that such a difference in trial duration does not affect memory scores. Box plots represent the median as the midline and the 25/ 75 % and 10/90 % quantiles as box boundaries and whiskers, respectively. Sample sizes are displayed within the Figure. # reflects significance relative to chance levels. * and ns above horizontal lines reflect significance, or lack thereof, in MWU-test or KW-tests. Statistical results are reported along with the source data in the supplemental data file “Thoener et al. DATA”. See **Fig. S2** for preference scores underlying the Memory Scores.

Given this separation of reward and punishment signalling at the level of the DANs, it was striking to observe that in both larval and adult *Drosophila* individual DANs can confer timing-dependent valence reversal. In the case of larvae, presentation of an odour followed by optogenetic activation of the DAN-i1 neuron (as covered by the SS00864-Gal4 strain, henceforth DAN-i1^864^) (forward training) leads to appetitive reward memory, whereas presentation of the odour upon termination of DAN-i1^864^ activation (backward training) establishes aversive frustration memory (Saumweber et al., 2018) (adults: Handler et al., 2019). Whether the mechanisms of frustration memory established by termination of DAN-i1^864^ activation differ from those of the likewise aversive memory established by unpaired presentations of odour and DAN-i1^864^ activation (Schleyer et al., 2018, 2020) remains to be tested. The DAN-f1 neuron, in contrast, supports aversive punishment memory upon forward training and appetitive ‘relief’ memory upon backward training (Weiglein et al., 2021; using SS02180-Gal4 as the driver, henceforth DAN-f1^2180^) (adults: Aso and Rubin, 2016; Handler et al., 2019; König et al., 2018). Whereas for DAN-f1^2180^ Weiglein et al. (2021) have characterized punishment and relief memories in some detail, no further analysis has been performed for the reward and frustration memories established by DAN-i1^864^ activation. Here, we undertake such an analysis to provide a more complete picture of the timing-dependent valence reversal conferred by dopaminergic neurons.

## Materials and Methods

### Animals

Experiments were performed on 3^rd^ instar foraging larvae of *Drosophila melanogaster* raised on standard food at 25 °C, 60-70 % relative humidity, in a room running on a 12 h light/dark cycle but in vials wrapped with black cardboard to keep them in darkness.

We used the split-Gal4 driver strain SS00864 (HHMI Janelia Research Campus, USA; Saumweber et al., 2018), supporting strong and reliable transgene expression in DAN-i1 of both hemispheres, plus expression from a few additional cells that is stochastic between hemispheres and preparations (Saumweber et al., 2018; Schleyer et al., 2020; Weiglein et al., 2019). As argued before, such stochastic expression is unlikely to cause systematic effects in the behavioural mass assays employed in the present study (Saumweber et al., 2018; Schleyer et al., 2020). We refer to this driver strain and the covered neurons as DAN-i1^864^.

For optogenetic activation, offspring of the UAS-ChR2-XXL effector strain (Bloomington Stock Centre no. 58374; Dawydow et al., 2014) crossed to DAN-i1^864^ were used. The driver control resulted from DAN-i1^864^ crossed to w^1118^ (Bloomington Stock Centre no. 3605, 5905, 6326), the effector control from UAS-CHR2-XXL crossed to a strain carrying both split-Gal4 landing sites (attP40/attP2) but without Gal4 domains inserted (“empty”) (HHMI Janelia Research Campus, USA; Pfeiffer et al., 2010).

### Odour-DAN associative learning

Procedures followed Saumweber et al. (2018) and Weiglein et al. (2021). Variations in the following procedures will be mentioned in the Results section.

Experiments were performed in a custom-made setup, consisting of a wooden box equipped with a light table featuring 24 x 12 LEDs (peak wavelength 470 nm; Solarox, Dessau-Roßlau, Germany) and a 6 mm thick diffusion plate of frosted acrylic glass on top to ensure uniform blue light for ChR2-XXL activation (120 μW/cm^2^). On top of the diffusion plate, Petri dishes were placed into a polyethylene diffusion ring illuminated by 30 infrared LEDs (850 nm; Solarox, Dessau-Roßlau, Germany). For video recording a camera (Basler acA204090umNIR; Basler, Ahrensburg, 196 Germany) equipped with an infrared-pass filter was placed approximately 25 cm above the Petri dish. The Petri dishes were filled with 1% agarose solution (electrophoresis grade; CAS: 9012-36-6, Roth, Karlsruhe, Germany) as the substrate on which cohorts of approximately 30 larvae were free to move once transferred from their food vials.

During training, one set of larvae was presented with an odour paired with optogenetic activation of DAN-i1^864^ at the mentioned inter-stimulus-interval (ISI), whereas a second set of larvae received the odour and the DAN-i1^864^ activation in an unpaired manner (**Fig. S1**). Two different ISIs were used: −10 s for forward training (the 30-s odour presentation preceded the 30-s DAN-i1^864^ activation by 10 s) and 30 s for backward training (the odour presentation occurred 30 s after DAN-i1^864^ activation had started) (**Fig. S1**).

For the presentation of the odour (*n*-amylacetate, AM; CAS: 628-63-7, Merck, Darmstadt, Germany, diluted 1:20 in paraffin oil; CAS: 8042-47-5, AppliChem, Darmstadt, Germany) or of paraffin oil as the solvent control (S), we used Petri dish lids equipped with four sticky filter papers onto which 5 μl of either substance could be applied. Paraffin oil does not have behavioural significance as an odour (Saumweber et al., 2011).

After three training trials, the larvae were placed in the middle of a fresh test Petri dish and given the choice between AM and S on opposite sides. After 3 min, the number of animals (#) on either side and in a 1-cm wide middle stripe was determined and the preference for the odour was calculated as:

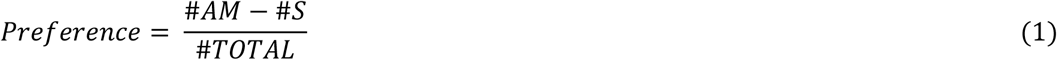

Thus, values can range between 1 and −1 with positive values indicating attraction and negative values aversion to AM.

From paired-trained versus unpaired-trained sets of larvae a Memory Score was calculated as:

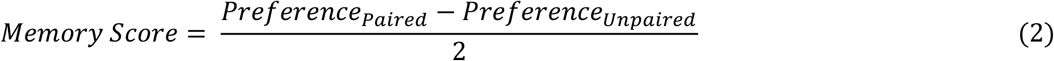

Thus, Memory Scores range between 1 and −1, with positive values indicating appetitive associative memory and negative values aversive associative memory. Each sample (N = 1) thus reflects the behaviour of two cohorts of approximately 30 larvae, one paired-trained and the other unpaired-trained.

### Video-tracking of locomotion

During the test, larval behaviour was video-recorded and analysed offline (Paisios et al., 2017). The typical zig-zagging larval behaviour was classified as either a head cast (HC) or a run, and was characterized by the HC rate, the change in orientation that results from a HC, and the run speed. All measurements are presented combined for around 30 larvae on a given Petri dish as one sample (N= 1).

The HC rate-modulation was defined as:

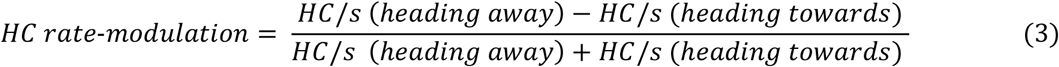

Positive scores indicate that larvae carry out more HCs per second (HC/s) while heading away from the odour than while heading towards it, and thus indicate attraction. In contrast, negative scores indicate aversion.

To calculate the difference in the HC rate-modulation of animals after paired versus unpaired training, we calculated the ΔHC rate-modulation as:

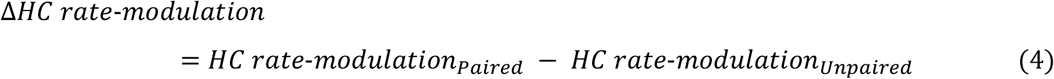

The Reorientation per HC was defined as:

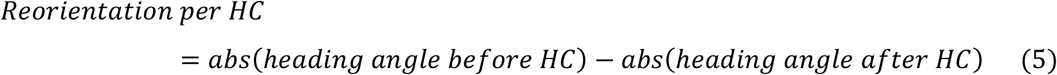

The absolute heading angle was defined as 0° when the animal’s head was pointing towards the odour and 180° when pointing away from it. HCs reorienting the animals towards the odour thus reduce this absolute heading angle and yield positive scores indicative of attraction whereas HCs reorienting the animal away from the odour result in negative scores indicative of aversion.

To calculate the difference in reorientation of animals after paired versus unpaired training, we calculated the ΔReorientation per HC as:

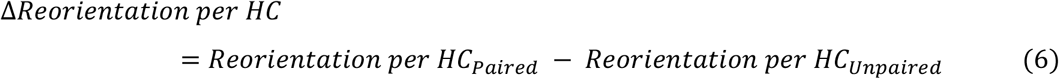

Run speed-modulation was defined as:

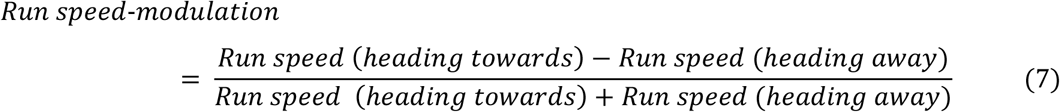

Thus, positive scores result if animals run faster when heading toward the odour than when heading away, indicating attraction, whereas negative scores imply aversion.

To calculate the difference in Run speed-modulation of animals after paired versus unpaired training, we calculated the ΔRun speed-modulation as:

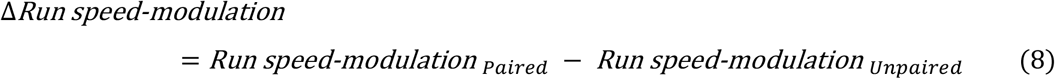

In all three cases, positive Δ scores would thus indicate appetitive memory. In turn, negative Δ scores would indicate aversive memory.

### Statistics

Non-parametric statistics were performed throughout (Statistica 13, RRID:SCR_014213, StatSoft Inc., Tulsa, USA). For comparisons with chance level (zero), one-sample sign tests (OSS) were used. To compare across multiple independent groups, we conducted Kruskal-Wallis tests (KW) with subsequent pair-wise comparisons by Mann-Whitney U-tests (MWU). To ensure a within-experiment error rate below 5% Bonferroni-Holm correction (Holm, 1979) was applied. Box plots show the median as the middle line, the 25 and 75 % quantiles as box boundaries, and the 10 and 90 % quantiles as whiskers. Sample sizes were chosen based on previous, similar studies. Experimenters were blind with respect to genotype when applicable. The results of the statistical tests and the source data of all experiments performed are documented in the Data file ‘Thoener et al. DATA’. Experimenters were blind to genotype.

## Results

We crossed the DAN-i1^864^ driver to the UAS-ChR2-XXL effector and trained larvae such that the odour was presented either 10 s before DAN-i1^864^ activation started (forward paired training) or 30 s after the start of DAN-i1^864^ activation (backward paired training; **Fig. S1**). To determine the memory score in each case the odour preference of the larvae was compared with the preference of larvae that had undergone presentations of the odour unpaired from DAN-i1^864^ activation (unpaired). Forward training induced appetitive, reward memory, which was not the case in the genetic controls (**Fig. 1B**) (Saumweber et al., 2018). In contrast, backward training resulted in aversive, frustration memory (**Fig. 1C**) (Saumweber et al., 2018). Given the reports of a punishing effect of light (von Essen et al., 2011), the data for the driver control suggest a moderate appetitive relief memory introduced by backward training with light; notably, such an effect would lead us to underestimate the aversive frustration memory in the experimental genotype (**Fig. 1C**).

To further analyse reward and frustration memory by DAN-i1^864^ activation, we next enquired into the impact of the duration of DAN-i1^864^ activation. Despite a trend, this manipulation did not affect reward memory upon forward training (**Fig. 1D**). After backward training, however, we observed that frustration memory increased with longer-lasting DAN-i1^864^ activation before odour presentation (**Fig. 1E**).

Second, given that a single training trial can be sufficient to establish memory for some natural tastant reinforcers (Weiglein et al., 2019), we tested for one-trial memory using DAN-i1^864^ activation. We observed reward memory after one-trial forward training with DAN-i1^864^ activation, but no frustration memory after one-trial backward training (**Fig. 1F-G**). Accordingly, when compared across experiments reward memory does not benefit from repeated training (**Fig. 1B**, **Fig. 1D** middle plot vs. **Fig. 1F**), whereas frustration memory does (**Fig. 1C**, **Fig. 1E** middle plot vs. **Fig. 1G).**

Third, we were interested in how temporally stable reward and frustration memories established by DAN-i1^864^ activation are. Reward memory lasted at least 40 min and indeed only tendentially decayed across this time period (**Fig. 1H**). In contrast, frustration memory was observed for only up to 10 min after training (**Fig. 1I**). We note that the experiments shown in Fig. 1H and Fig. 1I used a shortened protocol with reduced idle times before and after the actual training events, such that the total trial duration was 8 rather than 12 min. In a direct comparison, this difference in procedure was without effect (**Fig. 1J-K**).

Fourth, we wondered whether reward and frustration memories established via DAN-i1^864^ activation differ in their locomotor ‘footprint’; that is, associative memories can modulate both rate and direction of lateral head movements (head casts, HC) (Paisios et al., 2017; Thane et al., 2019; Toshima et al., 2019). In comparison to larvae that underwent unpaired training, forward training promoted head casts when crawling away, rather than when crawling towards the odour, whereas the opposite was the case upon backward training (**Fig. 2A**). Likewise, forward and backward training promoted head casts reorienting the larvae towards and away from the odour, respectively (**Fig. 2B**). In addition, upon forward training we observed that the runs towards the odour were faster than those away from it (**Fig. 2C**); this was surprising given that in 13 previous datasets tendencies for such run speed modulation, which can be recognized in about half of the cases, had never reached statistical significance (Paisios et al., 2017; Saumweber et al., 2018; Schleyer et al., 2015; Schleyer et al., 2020; Thane et al., 2019; Toshima et al., 2019; Weiglein et al., 2019; Weiglein et al., 2021). In any event, these analyses show that reward and frustration memories established by DAN-i1^864^ are behaviourally expressed through opposite modulations of the same aspects of locomotion.

**Fig. 2.**
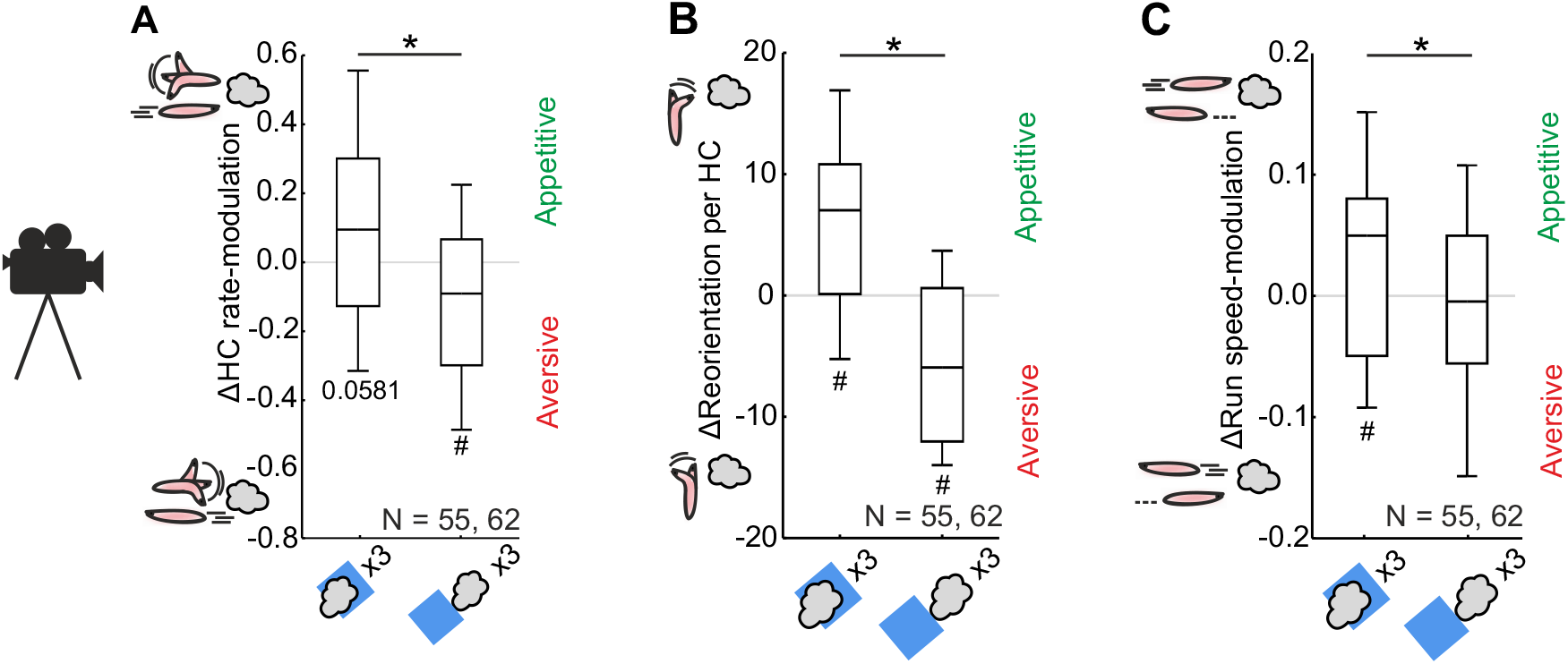
Locomotion footprint of reward and frustration memory by DAN-i1^864^ activation. Larvae were video-tracked during testing, and for three behavioural variables the difference between larvae undergoing forward- or backward-paired versus unpaired training (Δ) was calculated. **(A)** Forward training promoted head casts (HCs) when the larvae were crawling away from the odour rather than when crawling towards it. After backward training, the opposite was observed. **(B)** Forward and backward training prompted larvae to reorient their HCs towards and away from the odour, respectively. **(C)** Forward training resulted in faster runs towards the odour than away from it, whereas backward training had no effect upon run speed. Data are combined from Figures **1B-E** **(30-s light duration), J and K (12-min trial duration)**. Other details as in **Fig. 1**. See **Fig. S3** for results separated for forward- and backward-paired versus unpaired training. Statistical results are reported along with the source data in the supplemental data file “Thoener et al. DATA”.

## Discussion

We present a detailed characterization of the learning from the occurrence and termination of a central-brain reward signal, using optogenetic DAN-i1^864^ activation as a study case. Together with previous mirror-symmetric results concerning DAN-f1^2180^ (Weiglein et al., 2021), this reveals a 2×2 matrix of memory valence showing memories of opposite valence (appetitive or aversive) for stimuli that are associated with the occurrence or termination of central-brain reinforcement signals (**Fig. 3A**). Such a push-pull organization makes it possible to decipher the predictive, causal structure of events around a target occurrence and could be inspiring for computational modelling. Indeed, using notably broader drivers for effector expression, a similar organization was reported for adult flies (Handler et al., 2019) and may thus reflect a more general principle (Gerber et al., 2019). Elegant and general as such a 2×2 organization appears to be, there are a number of differences in the memories established:

- In both life stages, reinforcing effects were strongly determined by the time point of the occurrence/ termination of reinforcement, whereas its duration was only of impact for memories related to reinforcement termination (**Fig. 1B-E**; larvae DAN activation: Weiglein et al., 2021; adults DAN activation: König et al., 2018; electric shock: Diegelmann et al., 2013; Jacob and Waddell, 2020).
- Memories established through the occurrence of an event last longer than those established through its termination, in both larval and adult *Drosophila* (**Fig. 1H-J**; Weiglein et al., 2021; adults: Diegelmann et al., 2013; Yarali et al., 2008). With due caveats concerning rates of acquisition in mind, it might thus be that memories related to the occurrence of reinforcement are more stable than memories related to its termination.
- Fewer trials were sufficient to establish reward memory than frustration memory (**Fig. 1F-G**), which is in line with the opponent-process theory of Solomon and Corbit (1974). However, punishment and relief memory seem to benefit in a similar way from repeated training (larvae: Weiglein et al., 2021; adults: König et al., 2018).
- So far, only two out of three reinforcing DANs tested in larvae and two out of nine reinforcing DANs tested in adult *Drosophila* have been found to mediate opposing memories for stimuli associated with the occurrence versus termination of their activation (larvae: Saumweber et al., 2018; Weiglein et al., 2021; adults: Aso and Rubin, 2016; König et al., 2018). In other words, reinforcement signals that do *not* feature timing-dependent valence reversal need to be considered.

**Fig. 3.**
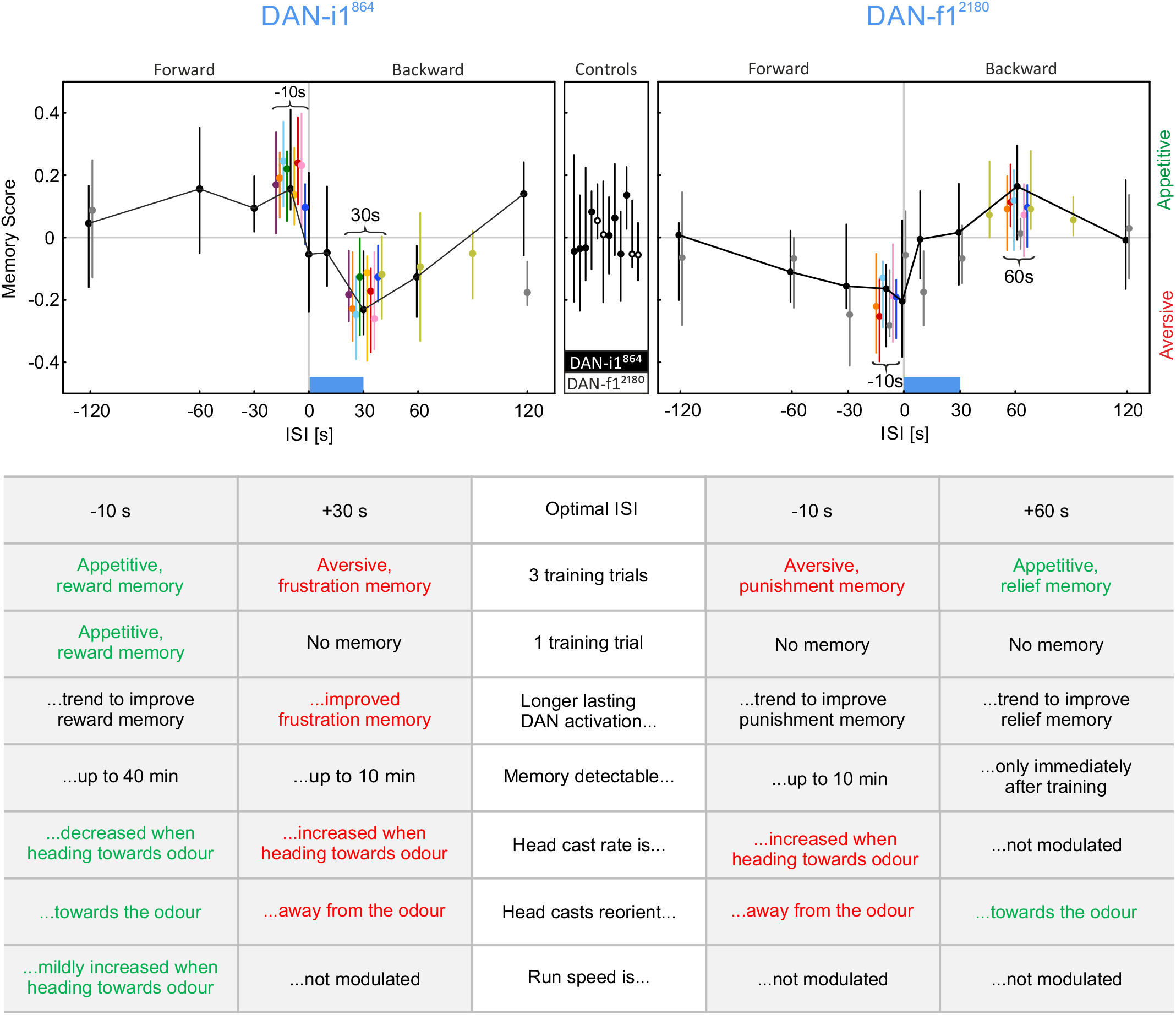
Summary matrix of timing-dependent valence reversal. Summary of the present and previously published data, showing a 2×2 matrix of timing and valence for DAN-i1^864^ (left panel) and DAN-f1^2180^ (right panel). Larvae underwent pairings of odour and optogenetic activation of the respective DAN (blue box) with different inter-stimulus intervals (ISIs). Odour presentation preceding DAN activation results in negative ISIs (forward training); the reversed sequence results in positive ISIs (backward training). The graph in the middle displays the results of the genetic controls. Shown are medians and 25/ 75 % quantiles of independent experiments from the present study and Saumweber et al. (2018) (top left) as well as Weiglein et al. (2021) (top right). Data and their original publication are documented in the supplemental data file “Thoener et al. DATA”. The table summarizes features of reward and frustration memory, and of punishment and relief memory established by pairing odour with the occurrence or termination of DAN-i1^864^ and DAN-f1^2180^.

As both DAN-i1 and DAN-f1 are dopaminergic (Eichler et al., 2017), it appears straightforward that reward and frustration learning as well as punishment and relief learning are mediated by dopamine. This is likely to be the case, given related results using broader drivers, at least for reward memory (Rohwedder et al., 2016; Thoener et al., 2021) and punishment memory (Selcho et al., 2009), and it is consistent with findings in adult *Drosophila* (Aso et al., 2019; Handler et al., 2019) and in mammals (Navratilova et al., 2015; but see König et al., 2018; Niens et al., 2017).

Taken together, our results complete the characterization of memories brought about by the timed activation of different larval DANs. The current data regarding DAN-i1^864^ activation and the data regarding DAN-f1^2180^ (Weiglein et al., 2021) provide a critical step to understanding the fundamental features of reinforcement processing, and pave the way both for an improved modelling of neural networks of reinforcement learning (Springer and Nawrot, 2021) and for further research into the underlying molecular mechanisms.

## Acknowledgements

Discussions with Christian König (LIN), Paul Stevenson (U Leipzig) and Philippe N. Tobler (U Zürich), as well as technical and experimental assistance by Anna Ciuraszkiewicz-Wojciech, Bettina Kracht, Thomas Niewalda, Lukas Riemen, Madeleine v. Maydell, Lukas Kuhn, Louisa Warzog and Irina Feldbrügge are gratefully acknowledged. We thank R.D.V. Glasgow (Zaragoza, Spain) for language editing.

## Author contributions

Conceptualization: J.T., A.W., B.G., M.S.; Methodology: J.T., A.W., M.S.; Formal analysis: J.T., M.S.; Investigation: J.T., A.W.; Data curation: J.T., M.S.; Writing - original draft: J.T., M.S., B.G.; Writing - review & editing: J.T., B.G., M.S., A.W.; Visualization: J.T., A.W.; Supervision: M.S., B.G.; Project administration: M.S., B.G.; Funding acquisition: B.G., M.S.

