## Supplementary Figures for "Optogenetically induced reward and ‘frustration’ memory in larval *Drosophila*"

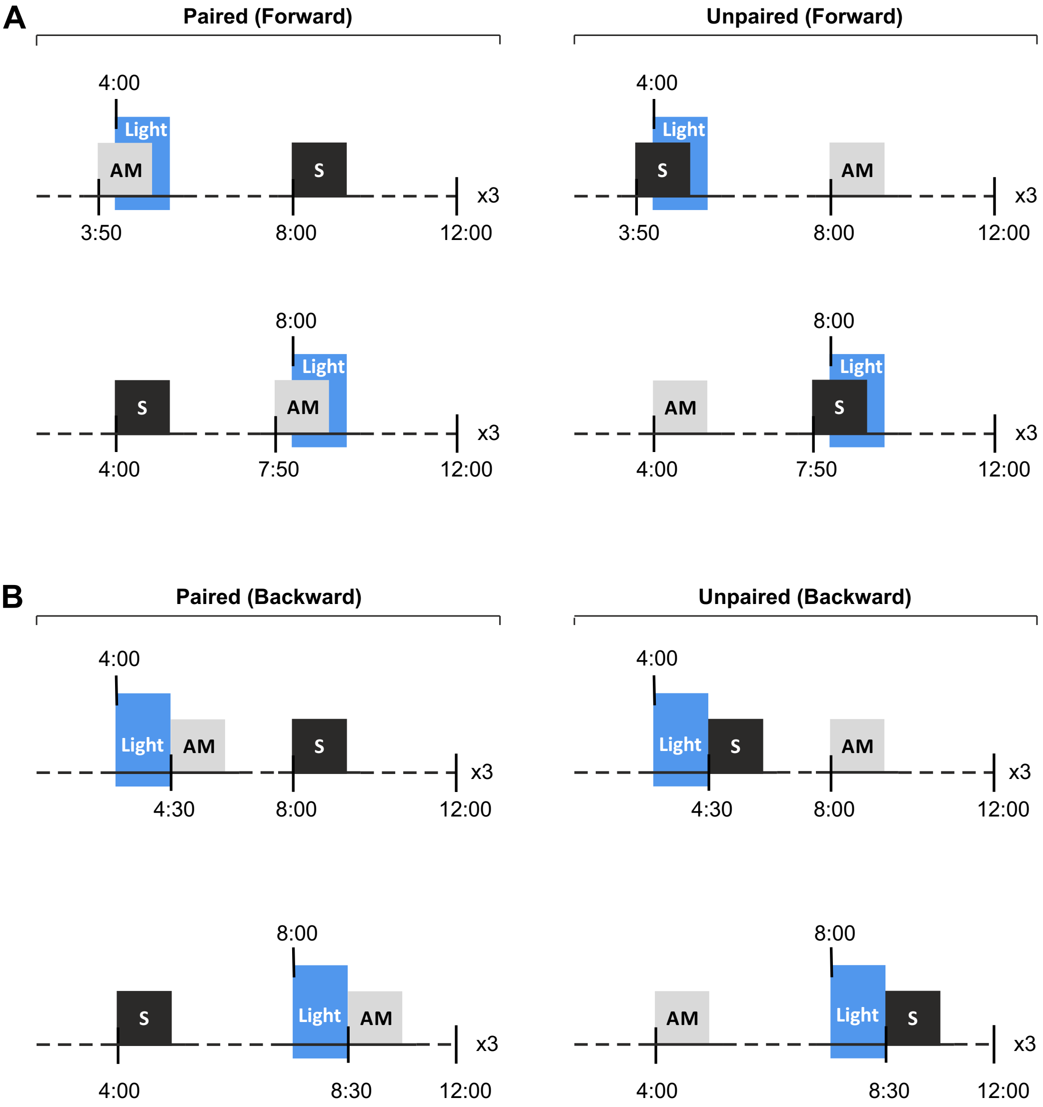


**Fig. S1.** Training procedure, showing that in the paired cohort, larvae were trained with timed presentation of odour (AM; grey box) together with optogenetic activation of DAN-i1$^{864}$ by blue light (Light; blue box), followed or preceded by presentation of the solvent as an 'odour control' (S; black box). Another cohort of larvae was trained with unpaired presentation of odour and light activation of DAN-i1$^{864}$. Both odour presentation and DAN-i1$^{864}$ activation by blue light lasted 30 s. In half of the cases, light activation started after 4 min, in the other half after 8 min. (A) Odour was either presented 10 s before optogenetic activation of DAN-i1$^{864}$ (forward conditioning; ISI -10 s) or (B) 30 s after blue light activation (backward conditioning; ISI +30 s). After three training trials, the preference for the odour was tested.

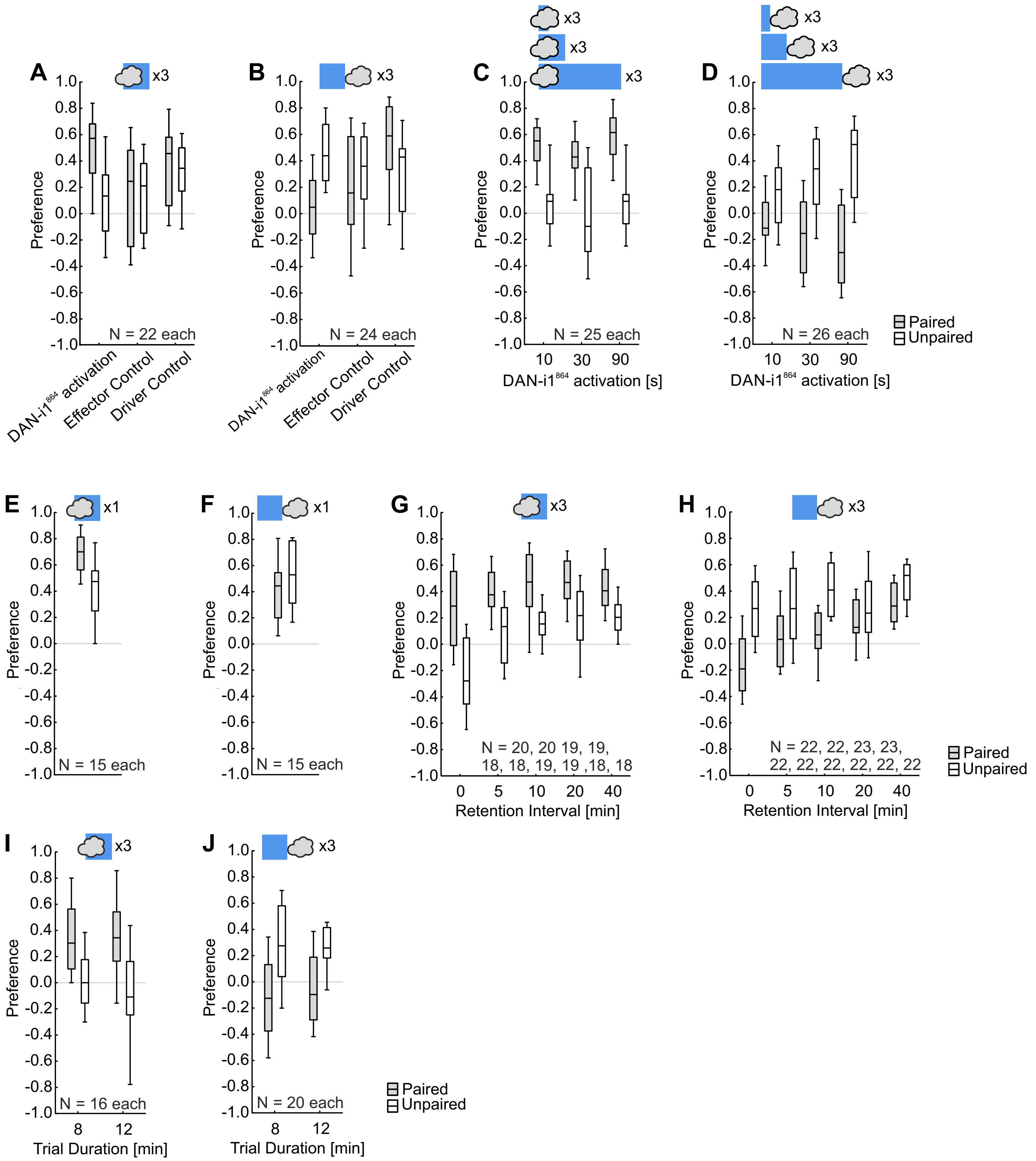


**Fig. S2.** Preference scores for the reciprocally trained sets of larvae underlying the Memory Scores from Fig. 1. Shown are Preference scores after paired (grey boxes) and unpaired (open boxes) training. Other details as in Fig. 1. Source data are documented in the supplemental data file “Thoener et al. DATA”.

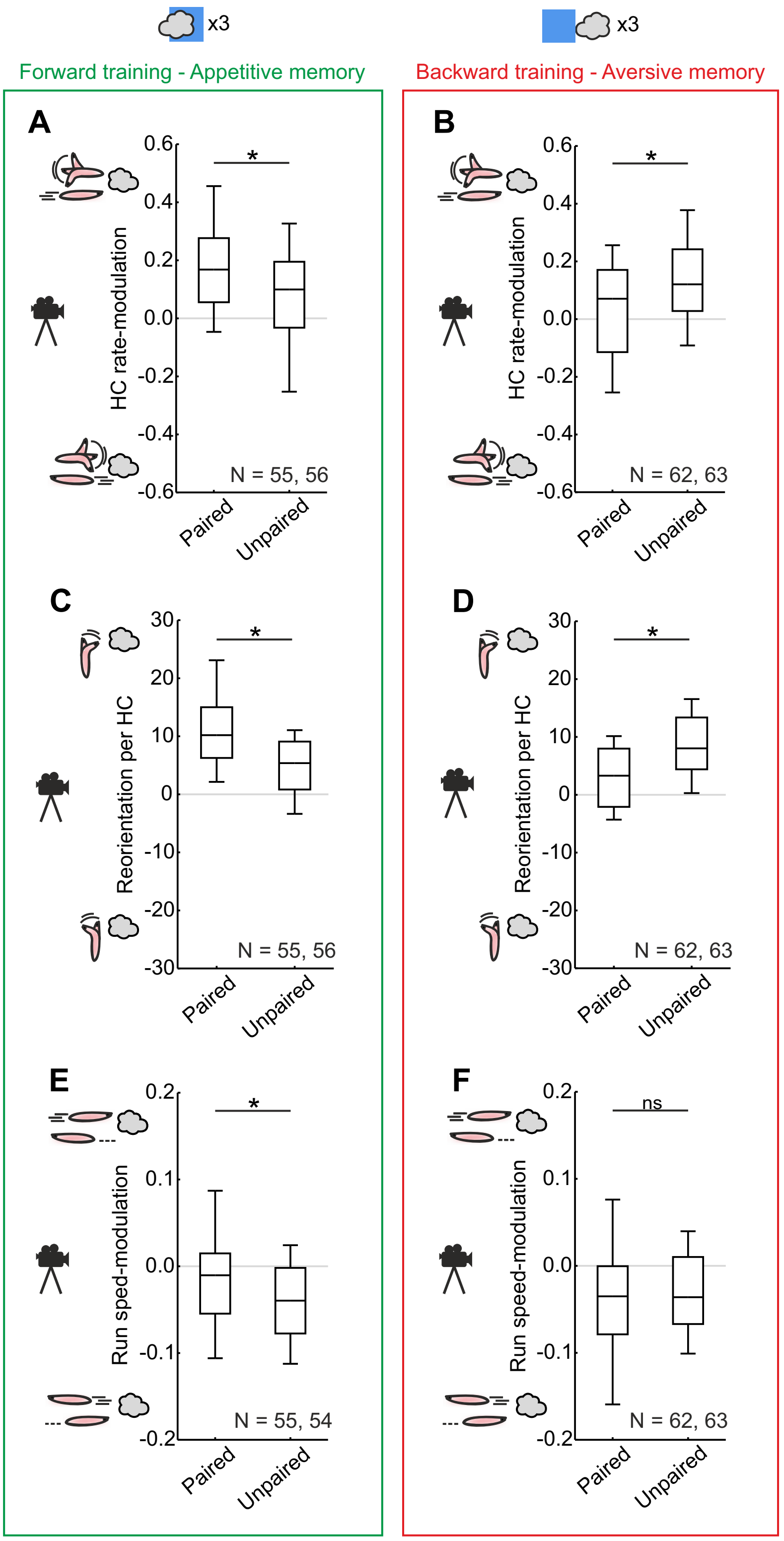


**Fig. S3.** Results separated for paired and unpaired training, underlying the D scores in Fig. 2. (A) Compared to larvae that were trained unpaired, after forward-paired training the larvae made fewer HCs when heading towards than heading away from the odour. (B) After backward training, the opposite was observed. (C) Larvae directed their HCs more towards the odour when they were trained forward-paired than when trained forward-unpaired. (D) After backward training, the opposite reorientation of the HCs was observed. (E) After forward-paired training, larvae run faster while heading towards the odour than while heading away, compared to larvae undergoing unpaired training. (F) No difference in run-speed was observed between backward-paired and unpaired-trained larvae. Data are combined from Fig. 1B-E (30-s light duration), J and K (12-min trial duration). Other details as in Fig. 1. Source data are documented in the supplemental data file “Thoener et al. DATA”.
